# Evolution of *sex hormone-binding globulin* gene expression in the primate testis

**DOI:** 10.1101/2024.08.05.606716

**Authors:** Warren M. Meyers, Geoffrey L. Hammond

## Abstract

In lower mammals testicular sex hormone-binding globulin (SHBG), also known as androgen binding protein, is well known to be a product of the Sertoli cells. However in humans, testicular SHBG is a product of the germ cells, is expressed from an upstream promoter and contains an alternative first exon 1A. Examination of testicular *SHBG* transcripts from members across primate suborders revealed that transcripts containing exon 1A are unique to Hominoids and Old World Monkeys. In contrast testicular *SHBG* transcripts in gray mouse lemur contained the proximal exon 1, while no evidence for *SHBG* expression could be detected in marmoset monkey testes. In general, the exonic identity of primate testicular *SHBG* transcripts could be predicted based on the structure of their gene’s 5’ regulatory region and we show that they change through the primate clade. This work provides insights into how molecular evolution of higher primate *SHBG* genes has resulted in distinct changes in how it is expressed in their testes.

## 1. Introduction

It has been long established that sex hormone-binding globulin (SHBG), also known as testicular androgen binding protein, ABP, is a product of the Sertoli cells in rodents ^1^. A protein species with similar physical and steroid-binding properties as SHBG in serum has also been demonstrated in human and primate testicular homogenates. However, its site of synthesis and cellular location was never confirmed and was always assumed to be a product of the Sertoli cells ^2–5^.

Expression of *Shbg* in rodent Sertoli cells originates from exon 1 and transcription from this region is under the control of the immediately flanking promoter (Figure 1). This was clearly demonstrated when rat *Shbg* transcripts were abundantly expressed within the Sertoli cells of a transgenic mouse containing a genomic fragment of the rat *Shbg* gene ^6^. In mice harbouring the corresponding 4.3 kb region of the human *SHBG* gene, no testicular *SHBG* transcripts were detected ^7^. Only when an 11 kb fragment of the human *SHBG* gene was inserted into the mouse genome were *SHBG* transcripts detected in their testes ^7^. Follow up reports indicated that they are a product of the germ cells and comprise an alternative first exon (Figure 1) ^8,9^. It was therefore clear that at least two molecular mechanisms were at simultaneously at work: (1) constitutive transcriptional repression of the human *SHBG* gene in Sertoli cells and (2) temporal activation of transcription of *SHBG* from an alternative promoter in testicular germ cells (Figure 1).

**Figure 1.**
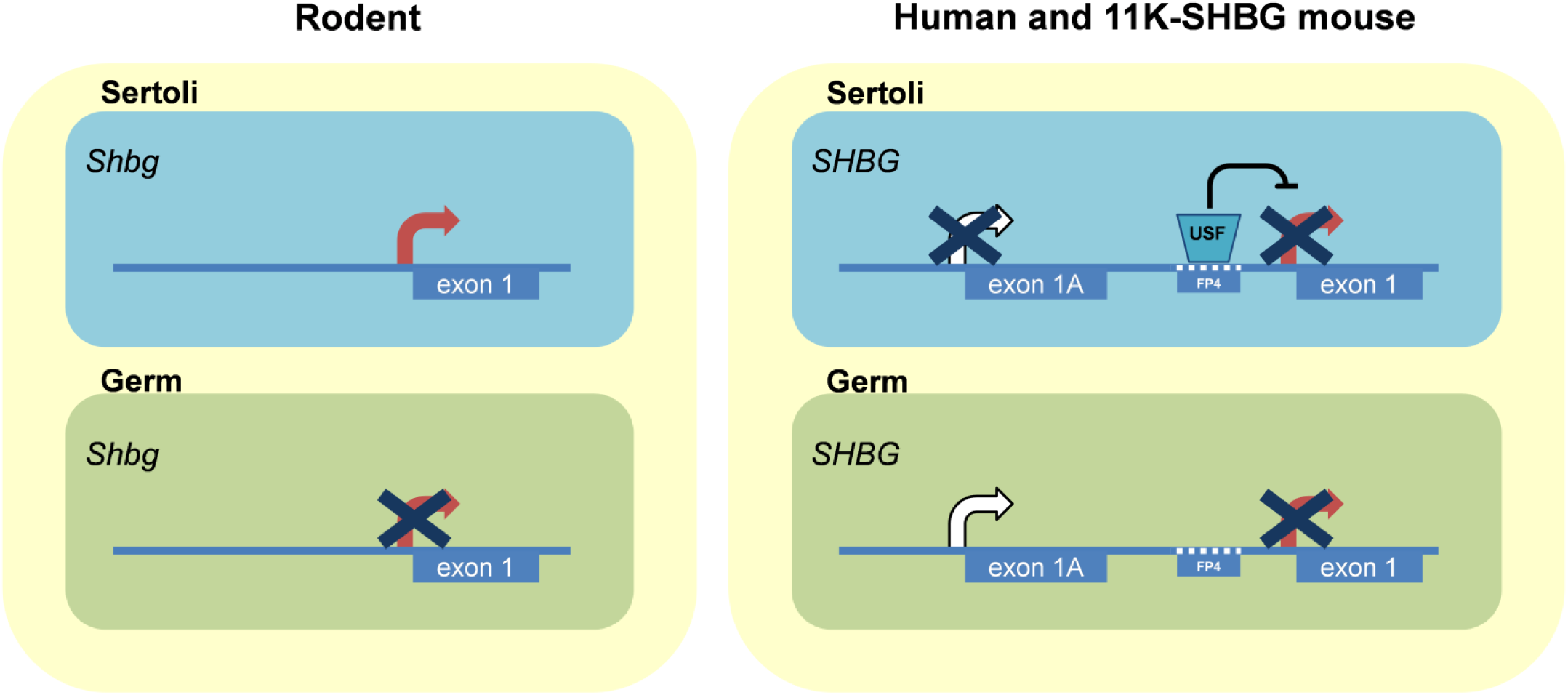
Distinct testicular cell-type expression profiles of *Shbg* and *SHBG* genes across mammals. In rodents and all lower eutherians known *Shbg* is expressed in their Sertoli cells from exon 1 ^1^. In contrast, in the human and 11K-SHBG transgenic mouse testis expression of *SHBG* occurs from an upstream alternative first exon in testicular germ cells ^8,11,17,45^. Expression of *SHBG* in Sertoli cells is constitutively repressed by USF1/2 from the FP4 element that is unique to humans and chimpanzees ^11^.

The genetic determinants that underlie the unique expression pattern of human *SHBG* in the testis have begun to be explored. Constitutive repression of *SHBG* in Sertoli cells is entirely dependent on the presence of a 38 bp element that was previously identified to be a DNase foot-printed region in hepatocytes (foot-print 4, FP4) ^10^. The centre of FP4 contains a binding site for upstream stimulatory factors (USF), and intriguingly, this element is the only major structural difference between the proximal promoters of the human and rodent *SHBG*/*Shbg* genes ^11^. When a line of transgenic mice carrying a modified human proximal *SHBG* transcription unit that lacked FP4 was derived, repression of *SHBG* was reversed and high levels of SHBG production resulted in their Sertoli cells ^11^.

The mechanisms underlying *SHBG* expression in testicular germ cells have received less attention. It is clear that the alternative transcription unit is regulated in a stage-dependent manner under the control of an upstream alternative promoter ^7,8^. In their phylogenetic analyses, Pinos and colleagues identified regions of near-perfect sequence identity to the human exon 1A within the *SHBG* genes of several higher primates ^12^. Their findings implied that these could be functional alternative first exons but analysis of testicular *SHBG* transcripts in these species is lacking. In addition to a putative exon 1A the chimpanzee *SHBG* gene also contains the FP4 element ^11^. Taken together this strongly suggests that *SHBG* expression in the chimpanzee testis is repressed in their Sertoli cells and is instead a product of the germ cells as in humans.

Considering the importance of these two genomic regions in directing the unique expression pattern of the human *SHBG* gene in the testis, we asked if they were common to all primates. With the aid of public nucleotide sequence databases and sequence alignment tools we constructed a phylogenetic comparison of *SHBG* 5’ regulatory regions using sequences from all major primate groups with the mouse *Shbg* gene as an outgroup. In particular, the regions responsible for repression of human *SHBG* in Sertoli cells and expression in germ cells were examined. This analysis is strengthened with RT-PCR studies for proximal and alternative *SHBG* transcripts using biopsies and cDNA obtained from several primate testes. We present novel data on the nature of testicular *SHBG* gene expression throughout the major primate groups and show that macromutation events in their *SHBG* genes are associated with changes in how it is expressed.

## 2. Materials and Methods

### 2.1 Nucleotide Sequences

The National Center for Biotechnology Information accession numbers for *SHBG/Shbg* sequences used in this study are as follows: *Homo sapiens*/ Human (NG_011981.1), *Pan troglodytes*/ Chimpanzee (NW_003458253.1), *Gorilla gorilla*/ Gorilla (NW_004006353.1), *Pongo abelii*/ Sumatran Orangutan (NW_002888527.1), *Nomascus leucogenys*/ Gibbons (NC_019834.1, NW_003501397.2), *Papio anubis*/ Olive Baboon (NC_018167.1, NW_003878344.1), *Macaca mulatta*/ Rhesus Macaque (NW_001102932.1), *Callithrix jacchus*/ Marmoset (ACFV01148737.1), *Saimiri boliviensis*/ Bolivian Squirrel Monkey (AGCE01058238), *Tarsius syrichta*/ Philippine Tarsier (ABRT02392649.1), *Microcebus murinus*/ Grey Mouse Lemur (ABDC01224946.1), *Mus musculus*/Mouse (AL731687.13).

### 2.2 RNA extraction and cDNA synthesis

Total RNA was extracted from tissues using the RNeasy Mini Kit (Qiagen) with a mid-protocol DNase I treatment. Concentrations of eluted RNA were determined using a NanoDrop 2000 spectrophotometer (ThermoFisher Scientific). Unless used immediately, RNA samples were stored at -80°C. Total RNA (2500 ng) was reverse transcribed using SuperScript® II Reverse Transcriptase with oligo(dT) primers (Thermo Fisher Scientific) in a 20 uL volume according to the manufacturer’s instructions with the exception that the synthesis incubation period at 42°C was increased to 60 min. All cDNA samples were stored at -20°C.

### 2.3 Primers and polymerase chain reaction (PCR)

Oligonucleotide/primer sequences for all PCR reactions described in this thesis are listed in Table 2.1. Amplification reactions were performed with either PCR Supermix (ThermoFisher Scientific) containing *Taq* polymerase, Phusion^TM^ High-Fidelity DNA polymerase for mutagenesis of C-G rich templates (ThermoFisher Scientific) or AccuPrime^TM^ *Pfx* DNA polymerase (ThermoFisher Scientific), each with their included reaction buffers. All PCR reactions were performed in 25 μL reaction volumes with primers diluted according to the instructions included with each polymerase. All reactions were carried out in a iCycler thermocycler (Bio-Rad) with the lid held at 100°C. Whole reaction volumes were analyzed by agarose gel electrophoresis stained with SYBR Safe DNA Gel Stain (ThermoFisher Scientific) diluted 1:10 000. Gel images were captured on a ImageQuant LAS 4000 system (GE Healthcare Life Sciences).

### 2.4 Primate genomic sequence retrieval and alignments

Primate *SHBG* sequences were retrieved from the NCBI public database online. For those sequences not annotated on GenBank, Nucleotide BLAST searches were performed using ∼300 bp regions of the human gene as query. Search sets were either reference genomic sequences (refseq_genomic) or whole-genome shotgun contigs (wgs) with the project or species/taxa of interest selected. Megablast, discontinuous megablast and blastn stringencies were used to locate sequences from species increasingly phylogenetically distant from human. NCBI accession codes for each species are listed in Table 1. Sequence alignments were performed using the Clustal Omega Multiple Sequence Alignment tool (http://www.ebi.ac.uk/Tools/msa/clustalo/). Percent sequence identity calculations were performed using the LALIGN server (http://www.ch.embnet.org/software/LALIGN_form.html) that applies the algorithm developed by Huang and Miller ^13^.

**Table 1.** Summary of *SHBG* genomic and testicular transcript features in relation to seminiferous tubule organization and testicular SHBG location in primates.

| Taxonomic group | *SHBG* gene features |  |  |  | Testicular *SHBG* transcripts |  | Known or inferred location of testicular SHBG | Seminiferous tubule organization [i] |
| --- | --- | --- | --- | --- | --- | --- | --- | --- |
|  | FP4/USF element | Sequence identity to human exon 1A (%) | Two intact C(A/G)CCC elements in alternative promoter | Other | Contains exon 1A | Contains exon 1 |  |  |
| Human | Yes |  | Yes |  | Yes [ii] | No [ii] | Germ cell [ii] | MS |
| Other Hominoids | Yes | 97 to 99 | Yes |  | nt | nt | *Germ cell* | MS |
| Old World Monkey | Yes | ~94 | Yes |  | Yes | No** | *Germ cell* | IM/MS |
| New World Monkey | Yes | ~90* | Yes | Exon 1A interrupted by *Alu* | No | No | *None?* | MS |
| Tarsier | No | 68.8 | No |  | nt | nt | *Sertoli cell* | Unknown |
| Lemur | No | 63.5 | No |  | No | Yes | *Sertoli cell* | SS |
| Mouse | No | 63.1 | No |  | No | Yes | Sertoli cell [iii] | SS |

[i] <sup>20, 44</sup>.
[ii] <sup>8,9</sup>.
[iii] <sup>46, 47</sup>.
\* Sequence identity not including *Alu* element.
\*\* Baboon testis contains *SHBG* transcripts with exon 1 at very low levels.
nt: not tested, MS: multistaged, IM: intermediate staged, SS: single staged.
Italicised entries indicate inferred or hypothesized location of testicular SHBG based on RT-PCR analyses and genomic trends described herein.

### 2.5 Non-human primate cDNA and tissue biopsies

Non-human primate testis cDNA samples were a generous gift from Dr. Rüdiger Behr at the Deutsches Primatenzentrum in Göttingen, Germany. Baboon (*Papio hamadryas*), Lion-tailed macaque (*Macaca silenus*), and Marmoset (*Callithrix jacchus*) cDNA were 100 ng/μL, while that from Rhesus macaque (*Macaca mulatta*) was 50 ng/μL. RNA samples used as templates for these cDNAs were DNaseI treated prior to being primed by oligo(dT). RNA*later*®-treated (ThermoFisher Scientific) liver and testis biopsies (0.22-0.25 g) from two grey mouse lemurs (*Microcebus murinus*) were provided by Dr. Erin Ehmke at the Duke Lemur Center in Durham, North Carolina, USA. From each, a 20-30 mg piece was excised and disrupted in 600 μL RLT buffer (Qiagen) supplemented with β-mercaptoethanol using a free-standing rotor-stator tissue homogenizer. Total RNA was extracted according to the RNeasy Mini Kit with a mid-protocol DNase I treatment.

## 3. Results

### 3.1 Alignment of primate *SHBG* 5’ regulatory regions

Building on earlier species comparisons ^11^ the region of the proximal human *SHBG* promoter that contains the FP4/USF element was aligned against the corresponding regions of ten other primate *SHBG* genes. Figure S1 clearly shows that while the FP4/USF element is present in all higher primate *SHBG* genes, it is absent in the tarsier and lemur sequences like the mouse. We then asked which primates had regions of their *SHBG* genes that resembled the human exon 1A and its flanking promoter. Alignment of the human alternative *SHBG* promoter and exon 1A with upstream regions of ten primate *SHBG* genes revealed that all simians contain regions with ≥90% sequence identity to the human exon 1A (Figures S2). The most striking finding of this alignment was that the exon 1A regions in both New World Monkey *SHBG* genes are interrupted by a 312 or 314 bp *Alu* element.

The presence of two CACCC elements within the alternative promoter region were also compared as we have recently shown they are required for human alternative *SHBG* promoter activity and are likely binding sites for KLF4^14^. While all species contained the proximal CACCC or functionally similar CGCCC element ^15,16^, the distal element is restricted to higher primates.

The sequence alignments in Figures S1 and S2 are summarized in Figure 2. All simiiforme *SHBG* genes contain the FP4 element which is sufficient criteria to suspect that *SHBG* is repressed in their Sertoli cells. Since exon 1A and its associated promoter in all OWM *SHBG* loci closely resemble those in the human, we hypothesized that these species express alternative *SHBG* transcripts in their testes and contain exon 1A. In the New World Monkey (NWM) the situation is less obvious. It is not clear if the presence of the *Alu* element interferes with transcription from this region since the exon 1A transcriptional start site and splice donor GT remain intact. Like the mouse *Shbg* gene, the lemur and tarsier lack the FP4 element. Their upstream regions corresponding to the human exon 1A have only 64-68% sequence identity. Based on their similarities to the mouse *Shbg* gene, we predict that lemur and tarsier testicular *SHBG* transcripts only contain exon 1.

**Figure 2.**
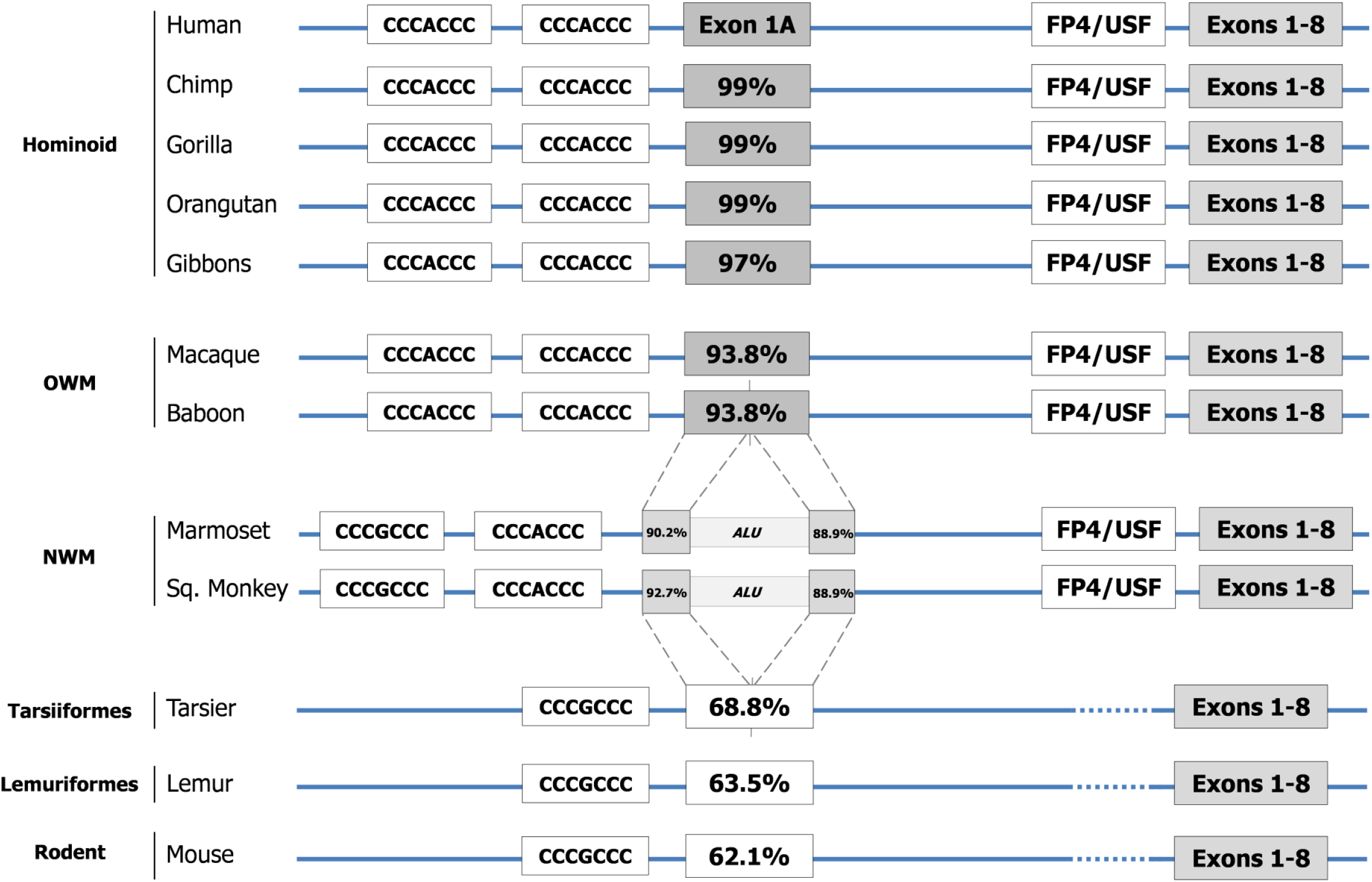
Summary of phylogenetic alignments of *SHBG* 5’ regulatory regions. All hominoids, OWM and NWM contain the FP4/USF element which represses *SHBG* expression in Sertoli cells^11^. OWM contain a region with very high (>90%) sequence identity to the human exon 1A. NWM also have a region that resembles exon 1A but it is interrupted by a 312/314 bp *Alu* element. The two C(A/G)CCC elements required for basal transcriptional activity from the alternative *SHBG* promoter are conserved in all hominoids, OWM and NWM.

To validate our conclusions from these sequence comparisons, we obtained whole-testis cDNA from several higher primates: cDNA from hamadryas baboon (*Papio hamadryas*), rhesus (*Macaca mulatta*) and lion-tailed (*Macaca silenus*) macaques represented Old World Monkeys while cDNA from three common marmosets (*Callithrix jacchus*) represented New World Monkeys. We also obtained liver and testicular biopsies from two grey mouse lemurs (*Microcebus murinus*) to represent the lower primates. With these samples RT-PCR analyses were performed for their *SHBG* transcripts.

#### 5.2.2 Old World Monkeys express alternative *SHBG* transcripts in their testes

Using primers with perfect complementarity to exons 1A, 1 and 8 on the baboon and macaque *SHBG* genes, we tested if OWM testes contain proximal or alternative *SHBG* transcripts. Figure 3 shows that both macaque and baboon testes are positive for *SHBG* transcripts containing exon 1A. The double banding pattern in the 1A-8 PCR assay resembles the doublet seen for *SHBG* transcripts in both human and 11K transgenic mice testes ^8,9^ where the upper band is the full length transcript and the lower lacks exon 7 ^8,9,17^. After 40 cycles of PCR both macaque testes were negative for proximal *SHBG* transcripts while the baboon testis produced a very low abundance of *SHBG* transcripts containing exon 1. Human hepatocellular carcinoma cell (HepG2) cDNA was used as a positive control for proximal (exons 1-8) *SHBG* transcripts.

**Figure 3.**
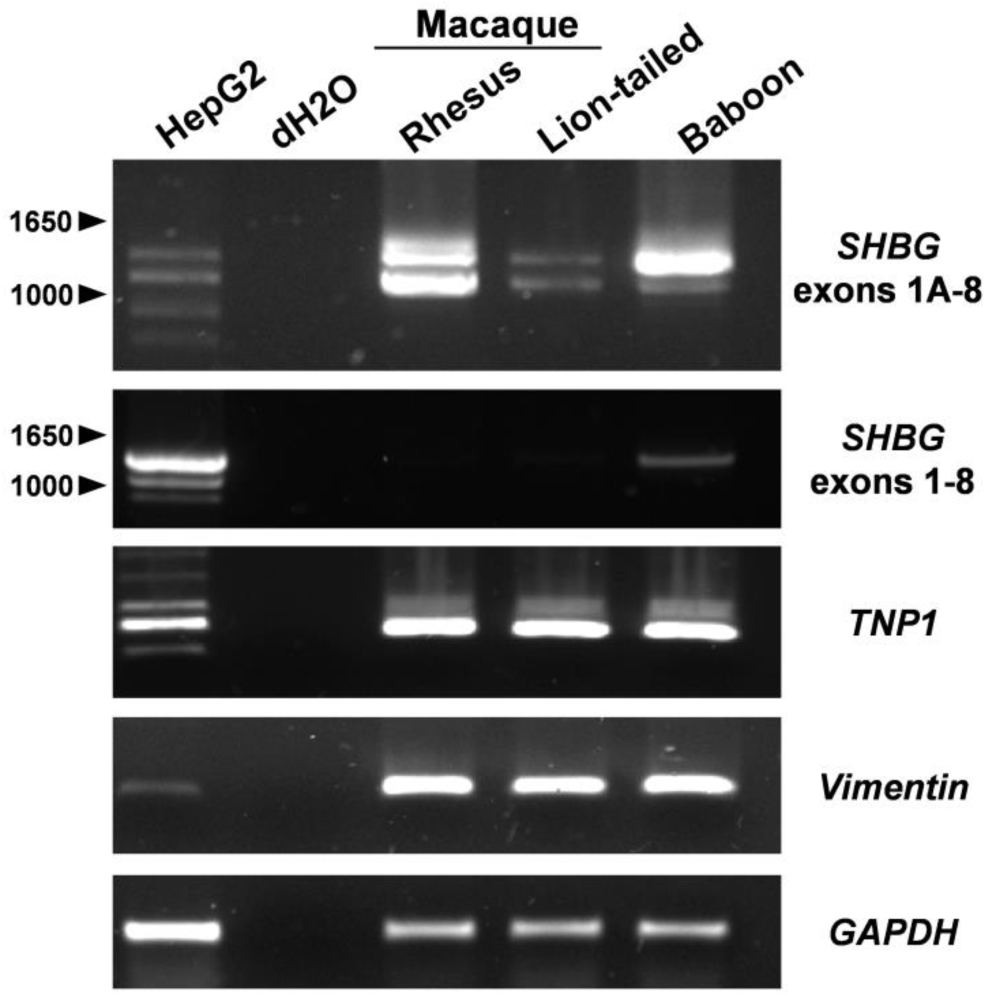
Alternative *SHBG* transcripts are present in Old World Monkey testes. RT-PCR assays for *SHBG* transcripts (exons 1A-8 and 1-8) were performed on cDNA derived from HepG2 cells, rhesus macaque, lion-tailed macaque and baboon testes. Analyses for *TNP1* and Vimentin transcripts indicate the presence of germ and Sertoli cells, respectively, in testis-derived cDNA. The integrity of all cDNA samples is assessed by analysis for *GAPDH* transcripts. All PCR products were subjected to electrophoresis on either 1% (for *SHBG*) or 1.5% (for *TNP1*, Vimentin and *GAPDH*) agarose gels. Molecular sizes in bp are indicated for the *SHBG* assays.

Furthermore, HepG2 cells have previously been shown to produce low levels of exon 1A-8 transcripts ^18^. Isolation and sequencing of the upper band from the rhesus 1A-8 assay revealed that it contains exon 1A followed by exons 2-8.

#### 5.2.3 No evidence for *SHBG* transcripts in the New World Monkey testis

Primers designed to marmoset *SHBG* exons 1A, 1 and 8 failed to detect alternative or proximal *SHBG* transcripts in the their testes. Since these PCR assays lacked an appropriate positive controls and do not rule out the possibility of a novel *SHBG* transcriptional start site used in their testis, primers were designed spanning the *SHBG* coding region (exons 3 and 8) with perfect complementarity to both the human and marmoset. Compared to the strong signal by HepG2 cDNA none could be detected in any of the marmoset testes despite their abundant signals for both transition protein 1 and vimentin transcripts indicating that these cDNA were derived from testes containing germ and Sertoli cells, respectively (Figure 4).

**Figure 4.**
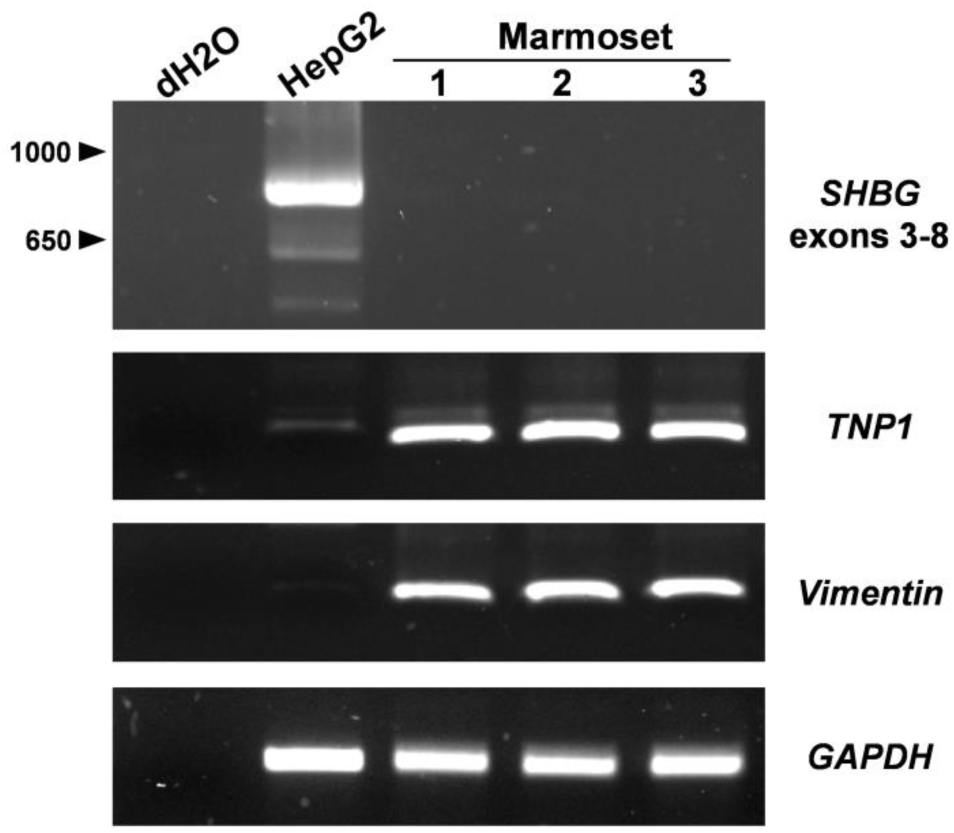
No evidence for *SHBG* transcripts in marmoset testes. RT-PCR assays for *SHBG* transcripts (exons 3-8) were performed on cDNA derived from HepG2 cells, and three marmoset testes. Analyses for *TNP1* and Vimentin transcripts indicate the presence of germ and Sertoli cells, respectively, in testis-derived cDNA. The integrity of all cDNA samples is assessed by analysis for *GAPDH* transcripts. All PCR products were subjected to electrophoresis on either 1% (for *SHBG*) or 1.5% (for *TNP1*, Vimentin and *GAPDH*) agarose gels. Molecular sizes in bp are indicated for the *SHBG* assay.

#### 5.2.4 High abundance of proximal *SHBG* transcripts in the lemur testis

Since the 5’ regulatory region of the lemur *SHBG* gene does not contain the FP4/USF repressor element, we expected that their testicular *SHBG* transcripts are produced from the same transcription unit used by the liver. Figure 5 shows that both lemur testes produce transcripts containing exon 1 as does the liver. Testing for alternative *SHBG* transcripts in the lemur testes was not performed because an appropriate positive control was lacking. A primer could not be designed with shared complementarity to the human exon 1A and the corresponding region of the lemur gene in order to use 11K-SHBG mouse testis cDNA as a control.

**Figure 5.**
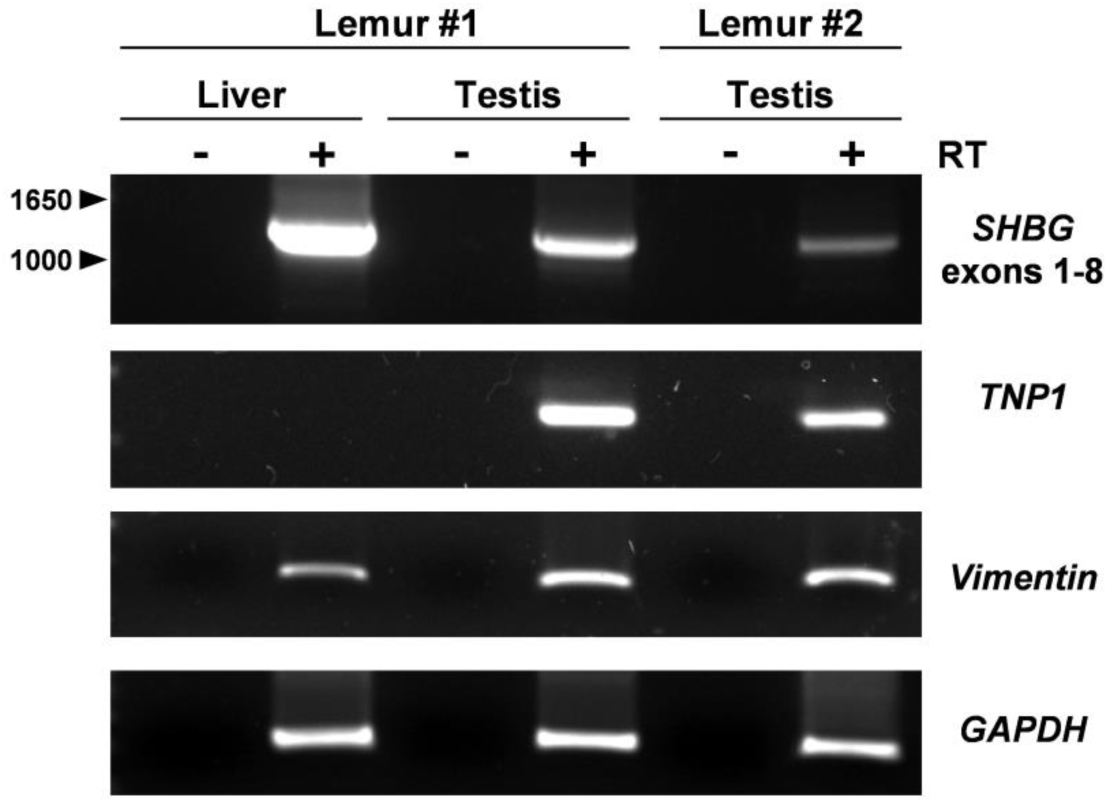
Lemur testes express *SHBG* transcripts from the proximal transcription unit. RT-PCR assays for *SHBG* transcripts (exons 3-8) were performed on cDNA derived from liver and testes from two grey mouse lemurs (*Microcebus murinus*). Analyses for *TNP1* and Vimentin transcripts indicate the presence of germ and Sertoli cells, respectively, in testis-derived cDNA. The integrity of all cDNA samples is assessed by analysis for *GAPDH* transcripts. All PCR products were subjected to electrophoresis on either 1% (for *SHBG*) or 1.5% (for *TNP1*, Vimentin and *GAPDH*) agarose gels. Molecular sizes in bp are indicated for the *SHBG* assay. RT, reverse transcriptase.

### 5.3 Discussion and future directions

Results from this study are summarized in Table 1. Primate *SHBG* genes containing the FP4/USF element are limited to all hominoids and OWM. On this criterion alone we hypothesized that these species do not contain *SHBG* transcripts containing exon 1 in their testes. This hypothesis was confirmed in macaques and marmosets. The baboon testis exhibited a very low abundance of exon 1-containing *SHBG* transcripts, however given the PCR product’s faint intensity compared to HepG2 cells after 40 amplification cycles, the presence of this transcript could represent transcriptional noise or degradation from sample quality. Follow up analyses for exon 1-containing *SHBG* transcripts are therefore warranted in other baboon testes. The lemur *SHBG* locus does not contain the FP4/USF repressor element. Consistent with this their testes are positive for transcripts arising from exon 1 indicating that the proximal transcription unit is a major source of SHBG in their testes. Since the 5’ regulatory region of the lemur *SHBG* locus closely resembles that on the mouse gene and that their testes utilize the proximal transcription unit it is very likely these transcripts are a product of their Sertoli cells just as in rodents.

All OWM *SHBG* genes contain regions with near-perfect sequence similarity to the human exon 1A. Since these species also have two intact CACCC activator elements it is likely that they all express alternative *SHBG* transcripts in their testes. All OWM samples tested contain an abundance of 1A-8 *SHBG* transcripts that are expected to be a product of their germ cells.

Failure to detect any *SHBG* transcripts in marmoset testes confirms that this gene is not expressed in this tissue from any transcription unit (Figure 4). Strikingly, this suggests that testicular SHBG is not required for marmoset fertility. Spermatogenesis in marmosets has many shared features with other higher primates, such as a multistaged seminiferous epithelium ^19–21^. In contrast, endocrine control of spermatogenesis as well as steroid hormone profiles in marmosets are much more distinct ^21^. Moreover, they also differ in other aspects of reproductive endocrine control. For instance, gonadotropin regulation of marmoset testicular steroidogenesis is mediated through chorionic gonadotropin (CG) instead of luteinizing hormone (LH) as in most other mammals ^22,23^. *CGβ* but not *LHβ* is expressed in the marmoset anterior pituitary ^24^. Exon 10 is spliced out of the marmoset luteinizing hormone receptor transcripts ^25^, resulting in a lost capacity to be stimulated by LH yet preserving stimulation capacity by CG ^26^.

All higher primate species studied to date, including marmosets, contain SHBG in their bloodstream ^27^. Marmoset SHBG is similar to human SHBG based on electrophoretic and relative steroid-binding properties ^28^. Pugeat et al ^29^ determined that plasma SHBG in NWM has lower affinity for testosterone (T) by about one order of magnitude as well as 3-6 fold higher levels compared to catarrhines. This may, in part, account for their 2-4 fold higher concentrations of free testosterone compared to other simians ^29^. The activity of 5α-reductase is drastically reduced in squirrel monkey androgen-sensitive tissues ^30^. This enzyme is required for the conversion of testosterone to 5α-dihydrotestosterone and reduced activity may help protect target-organs from excessive androgenization. NWM also exhibit an excess of total and free plasma glucocorticoid levels compared to catarrhines despite no physiological signs of glucocorticoid hormone excess ^31^. This is most likely due to a decreased affinity of the glucocorticoid receptor for its ligands ^31^ causing decreased negative feedback and increasing outflow from the hypothalamic-pituitary-adrenal axis. Interestingly, cloning and characterization of squirrel monkey corticosteroid binding globulin (CBG) revealed that it has a much lower binding affinity for glucocorticoids compared to human CBG ^32^. Increased circulating steroid hormone levels, decreased carrier protein affinity and either decreased nuclear receptor abundance or affinity to ligands are all features of the glucocorticoid, mineralocorticoid, sex steroid and vitamin D systems in NWM ^33^. It is not known what evolutionary pressures or advantages favoured their “generalized steroid hormone resistance” but is an important consideration when selecting NWM for studies relating to endocrine physiology.

Estimates of the diversification times of the major primate groups allows for approximation of when macromutation events occurred in *SHBG* loci during primate evolution. Since the FP4/USF element is common to all simiiformes it must have arisen in their common ancestor. Studies estimate that the simian-tarsiiforme divergence occurred either before 61-70 million years ago (Ma) ^34–36^ or before 81-82 Ma ^36–38^. Based on our knowledge that this element is sufficient to repress *SHBG* expression in Sertoli cells ^11^, its insertion into the genome of an early simian would have soon resulted in a phenotype of *SHBG* repression in the Sertoli cells of subsequent male offspring. The FP4 region is flanked by quadruplicate guanines (G_4_), and interestingly, there is a single G_4_ in the corresponding regions of the lower primates and mouse (Figure S1).

These repeat sequences allow us to hypothesize that the single G_4_ was an ancestral insertion site for the FP4/USF element within an early simian. This element is also remarkably small. Short interspersed nuclear elements (SINEs) are a type of small transposable elements on the order of 80-500 bp whereas other mobile elements range of one to many kb in length ^39,40^. Furthermore the origin of this element is unclear. A BLAST alignment of the FP4/USF element against the human genome returned no other matches other than itself in the *SHBG* gene, suggesting that it had not been copied from another part of the genome. No hits were also returned when it was aligned against the Philippine tarsier genome, suggesting that it is not a “jumping” mobile element that ancestrally existed at another site. Insertion of the FP4/USF element into the *SHBG* locus may therefore have arisen from some type of *de novo* virus-based or other unknown mechanism.

No evidence for *SHBG* expression in marmoset testes is attributed to exon 1A disruption by an *Alu*. This element is a feature in both squirrel monkeys and marmosets, two species of the *Cebidae* clade, one of the three major branches within NWM. Most analyses relying on fossil and molecular data suggest cebidae divergence from the rest of the NWM occurred before 20 Ma indicating that this insertion is at least that old ^36^. Loss of testicular *SHBG* expression can therefore only be concluded for squirrel monkeys and marmosets. Examination of *SHBG* loci for *Alu* insertion in exon 1A in the remaining NWM families (*Atelidae* and *Pitheciidae*) will allow me to determine if this is a feature of all platyrrhines. If they contain this *Alu* as well then loss of testicular *SHBG* expression can be estimated to be much older; before 35-45 Ma when platyrrhines diverged from catarrhines ^36^. Compared to other primates, marmosets have the highest incidence of *Alu* elements per megabase in their genomes (188 *Alu* counts/Mb), whereas humans and lemurs have 104 and 55 *Alu* counts/Mb respectively ^41^. The widely accepted mechanism for *Alu* insertion first involves transcription from another genomic *Alu*; the resulting transcript seeks out a nicked poly-thymine which is used to prime genomic extension along the *Alu* transcript ^42,43^. The insertion site utilized within NWM exon 1A is the 5’-TTTTTAAA-3’ that immediately precedes the *Alu* (Fig S2). Figure 2 indicates the high (∼90%) sequence identity of NWM exon 1A regions on either side of the *Alu*, therefore it is likely that before disruption this was a functional alternative first exon.

Another interesting observation from these data is that the presence of the FP4/USF element (and hence repression of Sertoli cell *SHBG*) is associated with primate species that exhibit a multi-staged arrangement of their seminiferous tubules ^20^. A given cross-section through a simiiforme testis will reveal >1 spermatogenic stage per seminiferous tubule, whereas lemur or rodent testes will show only a single-staged arrangement ^44^. Tarsiiformes are the only infraorder in the haplorhines whose seminiferous tubule arrangement has not been characterized ^20,44^ so it will be interesting to learn whether their tubules resemble those in lemurs or the rest of the haplorhines.

In this study, testicular transcript analyses were performed on heterogeneous cell populations containing both Sertoli and germ cells. In general, testis expression profiles of *SHBG* could be predicted based on the structure of that specie’s *SHBG* 5’ regulatory region. Based on our pre-existing knowledge of where *SHBG* and *Shbg* are produced in mammalian testes and the genomic and transcriptional trends from this study, we therefore infer that (1) expression of *SHBG* in testicular germ cells from an alternative first exon is a feature of all OWM and (2) both lemurs and tarsiers express *SHBG* in their Sertoli cells from exon 1 (Figure 6).

**Figure 6.**
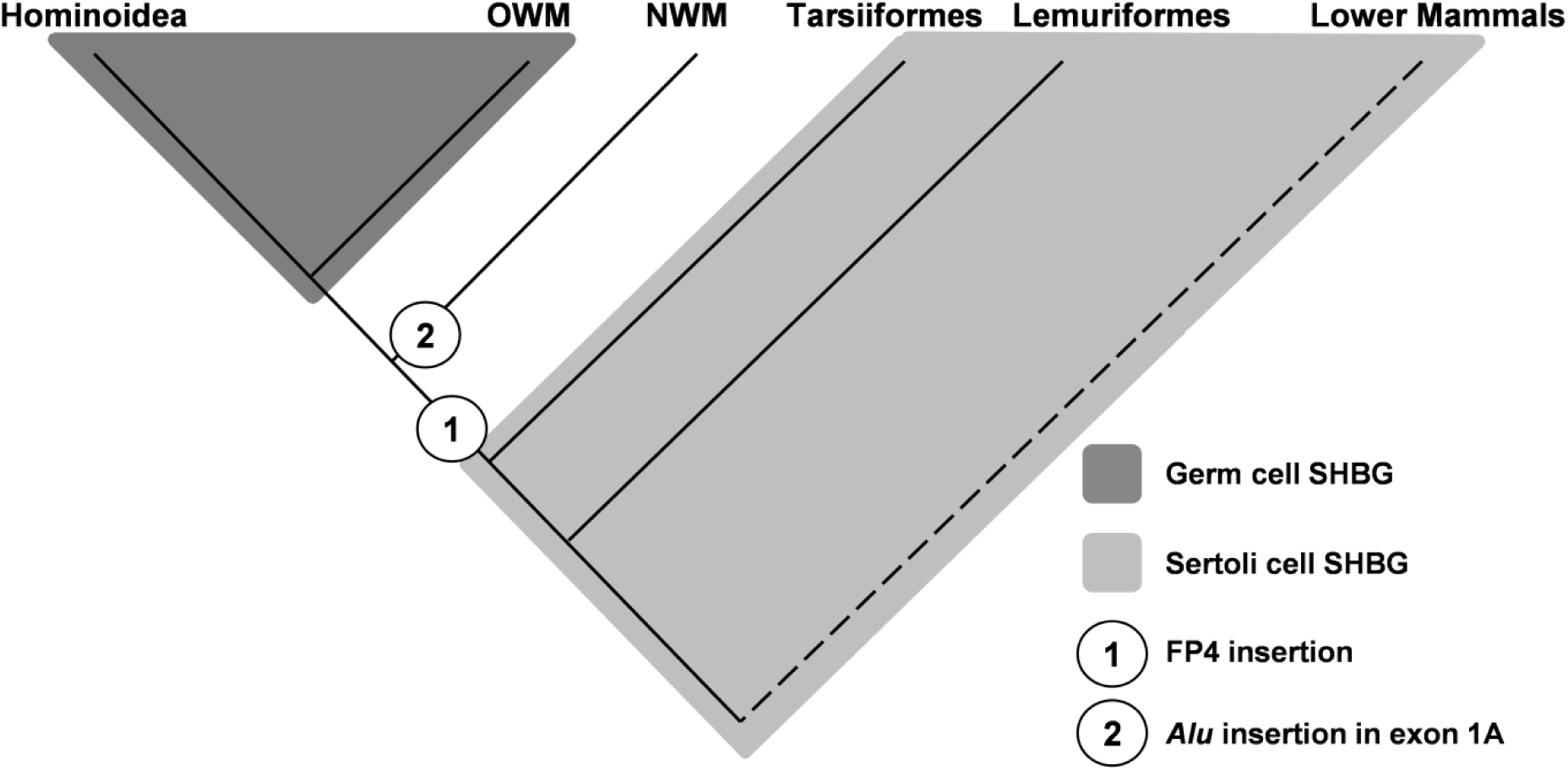
Trends of testicular *SHBG* gene expression across primates. Sertoli cells are the ancestral location of testicular SHBG. Alternative *SHBG* transcripts containing exon 1A have only been detected in hominoids and OWM (catarrhini) and are therefore the only groups where germ cell SHBG is expected to be synthesized. NWM do not express *SHBG* in their testes but probably did before *Alu* disruption on exon 1A. Each change in how testicular *SHBG* is expressed is always preceded by a structural change in the *SHBG* 5’ regulatory region of that group. Numbers indicate approximate period of a macromutation event in the *SHBG* gene: 1, insertion of the FP4 element. 2, insertion of an *Alu* element in the NWM alternative first exon. OWM, Old World Monkey, NWM, New World Monkey.

The origin of exon 1A is unclear. This transcription unit probably arose from molecular evolution of exon 1A so that it gained a splice donor site and its flanking promoter. Selection pressures in favour of germ cell SHBG are difficult to hypothesize, however its correlation with multistaged seminiferous tubules in simiiformes indicates that there were significant genetic and morphological changes in testis biology occurring around the same time during primate evolution. Knowledge on the arrangement of tarsier seminiferous tubules is absent in the literature and will be needed to test the strength of this association.

## Acknowledgements

We gratefully acknowledge the assistance of Dr. Erin Ehmke at the Duke Lemur Centre for providing us with lemur tissue biopsies as well as Dr. Rüdiger Behr at the Deutches Primatenzentrum for providing us with non-human primate testis cDNA. We also thank Drs. Sally Otto, Darren Irwin, as well as Chris Stinson at the Beaty Biodiversity Museum for use of their Certificate of Scientific Exchange under the Convention on International Trade in Endangered Species of Wild Fauna and Flora.

---

**Figure S1.**
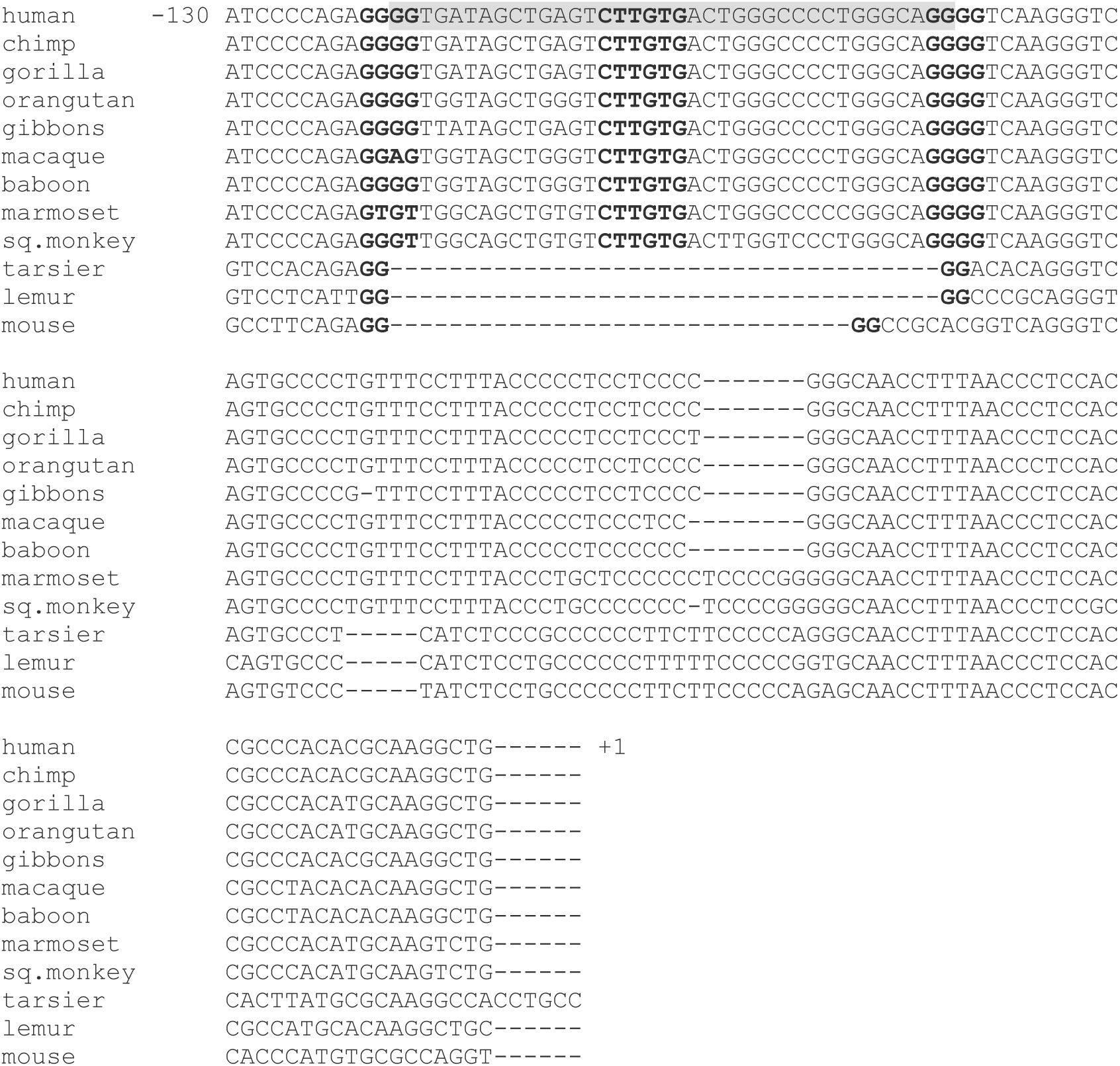
The FP4/USF element is limited to simiiforme primates. A minimal region of the human *SHBG* proximal promoter, from -130/+1 with respect to the transcriptional start site used by the liver ^10^ is aligned with the corresponding regions from the chimpanzee, gorilla, orangutan, gibbons, macaque, baboon, marmoset, squirrel monkey, tarsier, lemur and mouse *SHBG*/*Shbg* genes (Table 5.1). The region containing the entire FP4 element in the human sequence is shaded in grey. The central binding site for USF1/2 is bolded for emphasis ^11^ as are the quadruplicate guanines that form element’s boundaries.

**Figure S2.**
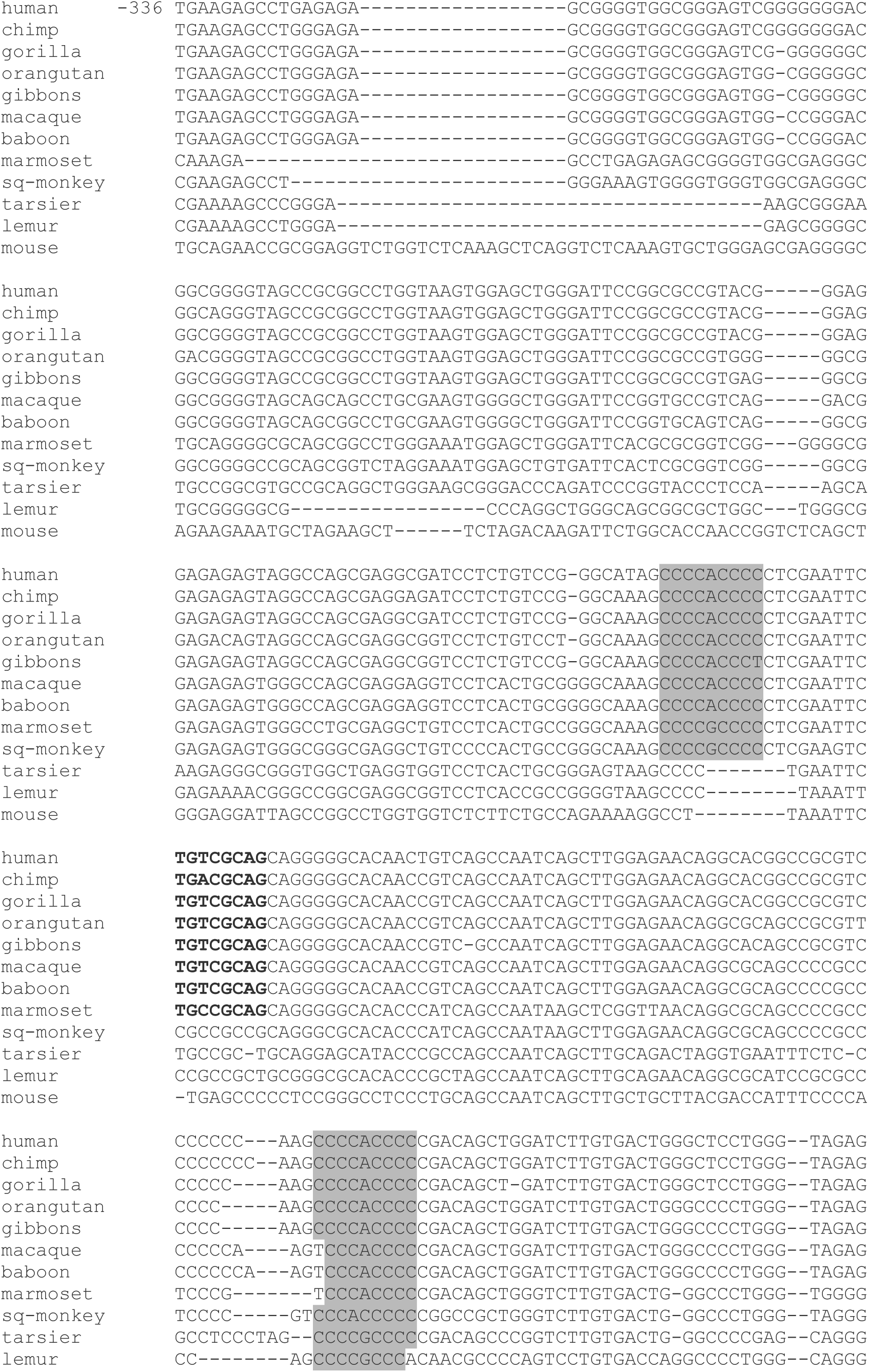
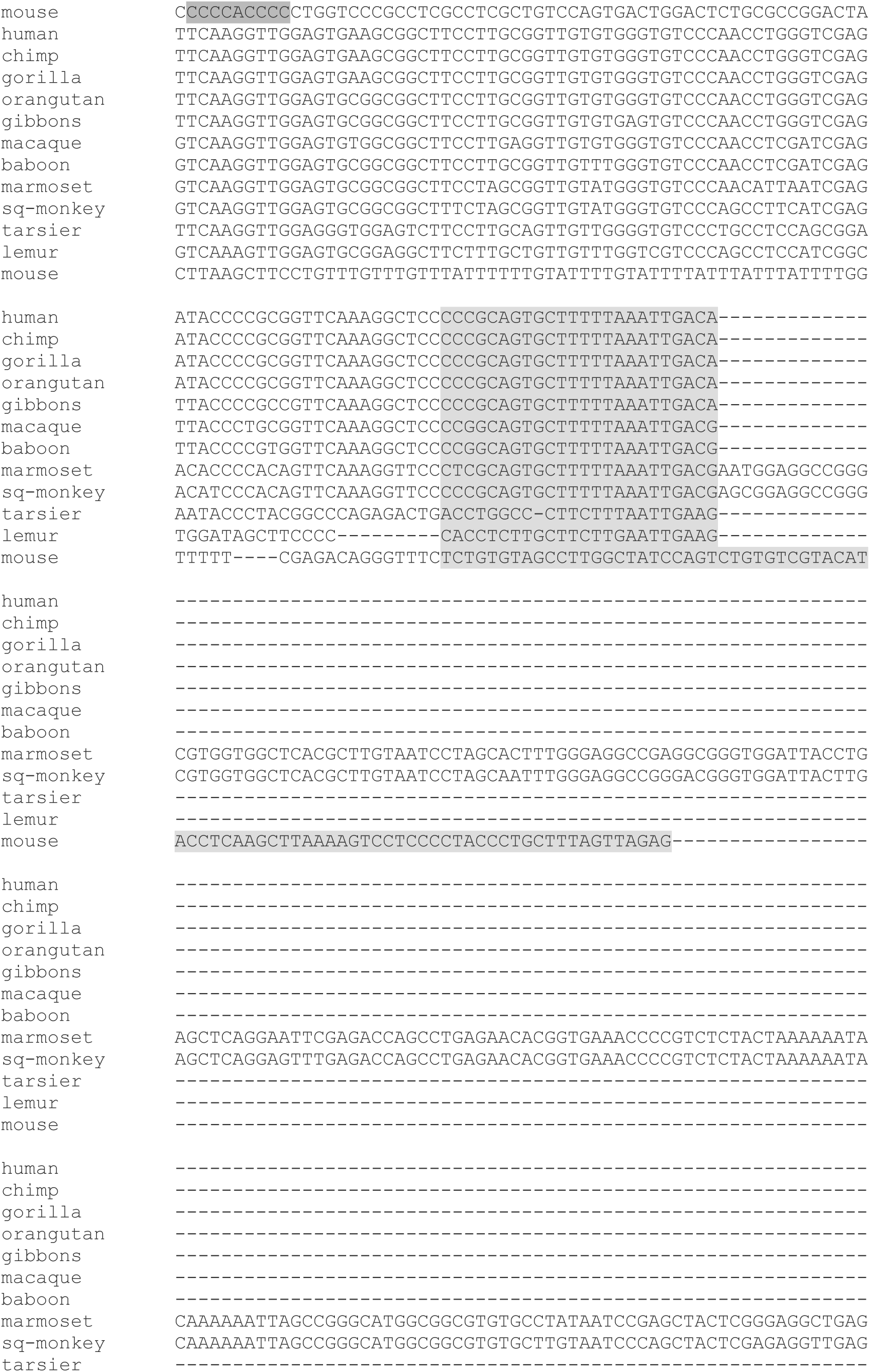
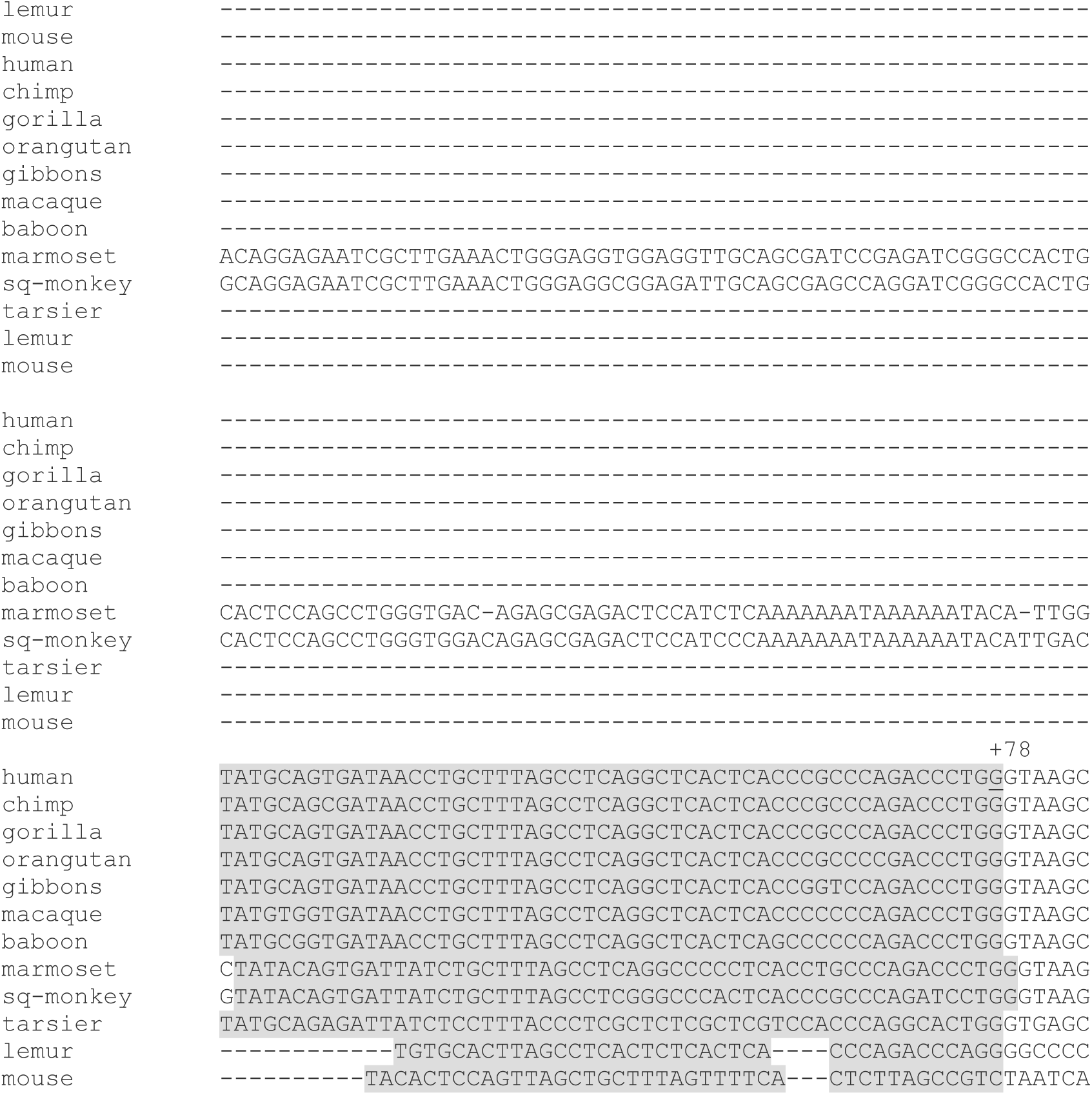
Comparison of the *SHBG* exon 1A and flanking promoter sequences among primates. The human alternative first exon and 336 bp of its 5’ flanking promoter used in testicular germ cells is aligned with the corresponding regions of the chimpanzee, gorilla, orangutan, gibbons, macaque, baboon, marmoset, squirrel monkey, tarsier, lemur and mouse *SHBG*/*Shbg* genes (Table 5.1). The marmoset and squirrel monkey exon 1A regions are each interrupted by an *Alu* element (312 and 314 bp, respectively). Lightly shaded bases indicate the human exon 1A and possible alternative first exons in other species. These regions were used to calculate the percent sequence identities listed in Figure 2 and Table 1. Intensely shaded bases indicate the CACCC regulatory elements described previously^14^. Bolded bases indicate a partial cyclic-AMP response element. The underlined guanine indicates the +78 position and 3’ boundary of the human alternative exon 1A.

